# Single-cell measurement of microbial growth rate with Raman microspectroscopy

**DOI:** 10.1101/2023.12.16.571966

**Authors:** Tristan A. Caro, Srishti Kashyap, George Brown, Claudia Chen, Sebastian H. Kopf, Alexis S. Templeton

## Abstract

Rates of microbial activity and growth are fundamental to understanding environmental geochemistry and ecology. However, measuring the heterogeneity of microbial activity at the single-cell level, especially within complex populations and environmental matrices, remains a forefront challenge. Stable Isotope Probing (SIP) is a standard method for assessing microbial activity and involves measuring the incorporation of an isotopically labeled compound into microbial biomass. Here, we assess the utility of Raman microspectroscopy as a SIP technique, specifically focusing on the measurement of deuterium (^2^H), a tracer of microbial biomass production. We generate calibrations of microbial biomass ^2^H values and find that Raman microspectroscopy reliably quantifies ^2^H incorporation ranging between 0 and 40 at. %. Applying the results of this calibration to a SIP model, we explicitly parameterize the factors controlling microbial growth quantification, demonstrating how Raman-SIP can measure the growth of microorganisms with doubling times ranging from hours to years. Furthermore, we correlatively compare our Raman-derived measurements with those of nanoscale secondary ion mass spectrometry (nanoSIMS) to compare the relative strengths of nanoSIMS- and Raman-based SIP approaches. We find that Raman microspectroscopy is a robust, accessible methodology that can readily differentiate and quantify the growth of individual microbial cells in complex samples.

**Importance:** Growth rate, the rate at which organisms grow and reproduce, is a key metric with which to evaluate microbial physiology and contributions to system-level processes. The heterogeneity of microbial growth across space, time, and populations is often difficult to capture with bulk-scale techniques. Single-cell methods hold promise for measuring the heterogeneity of microbial growth rates and responses to changing conditions *in situ*, without the need for cultivation of microbial isolates. In this study, we evaluated the ability of Raman microspectroscopy, a non-destructive and rapid technique, to measure the assimilation of isotopically labeled water into individual microbial cells and thereby calculate their rates of growth. We explicitly parameterize the factors controlling the quantification of microbial growth rate and compare this technique to standard methods. The framework we report allows researchers to couple single-cell and aggregate rate measurements to functional or system-level properties, a forefront challenge in microbiology.

## Introduction

Microbial growth rate is a critical parameter for assessing biogeochemical cycling, environmental habitability, and microbial fitness. Not only does the rate of microbial growth represent a geochemical transformation, but it is also a parameter that microorganisms modulate in response to changing conditions: microorganisms may more readily replicate when conditions are favorable to them and adopt alternative physiologic states under adverse conditions.

Methods for measuring microbial growth often rely upon the addition of an isotopically labeled substrate to a natural sample and measuring its incorporation into microbial biomass, a method generally termed stable isotope probing (SIP) (1–3). SIP has been applied with the stable isotopes of various elements including H, C, N, O, S, etc., through measurement of single-cells (4–8), membrane lipids (3,9,10), and nucleic acids (2,11–13). One approach to probe cellular biosynthesis is to amend an environmental sample with deuterated water (^2^H_2_O) (3–6,8,10,14–18). Water is incorporated into microbial biomass during the synthesis of lipids, nucleic acids, and proteins, and so probing a sample with deuterated water allows the quantification of microbial biomass growth. ^2^H_2_O has been termed a “passive” tracer for microbial anabolism, as it is nutritionally neutral and taxonomically agnostic.

In microbial communities, spatial relationships dictate ecological and evolutionary dynamics, as well as community structure and function (26–30). However, microbial growth is known to be extremely heterogenous in environments such as soil matrices (30), marine particles, (31–33), the rock-hosted subsurface (34,35) and others. Spatially-resolved measurements of microbial growth can identify the habitability and biological activity of environmental samples at the microscale. Furthermore, single-cell measurements of microbial growth are especially useful for microbial ecologists as they provide information on anabolic heterogeneity across and within populations. However, describing the spatial distributions of microorganisms in the environment, in addition to their degrees of activity, remains one of the primary challenges in microbial ecology.

Nanoscale secondary ion mass spectrometry (nanoSIMS) is a standard method for conducting spatially-resolved SIP measurements of individual cells (4,6,7,10,15,18,36–39). NanoSIMS relies on the measurement of secondary ions released via ablation by a primary ion beam and measures isotopes of multiple elements at high spatial resolution. However, isotope ratios measured via nanoSIMS may become depressed due to the inclusion of stains, nucleic acid probes, and sample washing (40–42). Additionally, nanoSIMS remains a costly and time intensive method that is inaccessible to many researchers. We therefore set out to assess alternative methodologies that may circumvent these difficulties.

In 2015, foundational work by Berry and colleagues established Raman microspectroscopy as a viable tool for detecting deuterium enrichment of microbial biomass at the single-cell level (8). Raman-based measurements of deuterium in microbial biomass rely on the principle that there exists a strong band centered at 2800 cm^-1^, corresponding to C-H bonding environments associated with lipids, proteins, and nucleic acids of cells. When deuterium is incorporated into microbial biomass, this band, now representing C-^2^H (“C-D”) bonds, becomes red-shifted and centers at 2200 cm^-1^ (**Fig. 1B**). In previous work, the “CD% metric”, defined as the fractional abundance of CD and CH peak areas, (CD% = CD / (CD + CH) × 100%) was found to correlate with the deuterium fractional abundance (*^2^F*) of the growth medium provided to the organisms (8). Berry et al. illustrated applications of Raman-SIP in combination with fluorescence in situ hybridization (FISH) and cell-sorting methodologies to capture the identity of active microbial community members.

**Figure 1:**
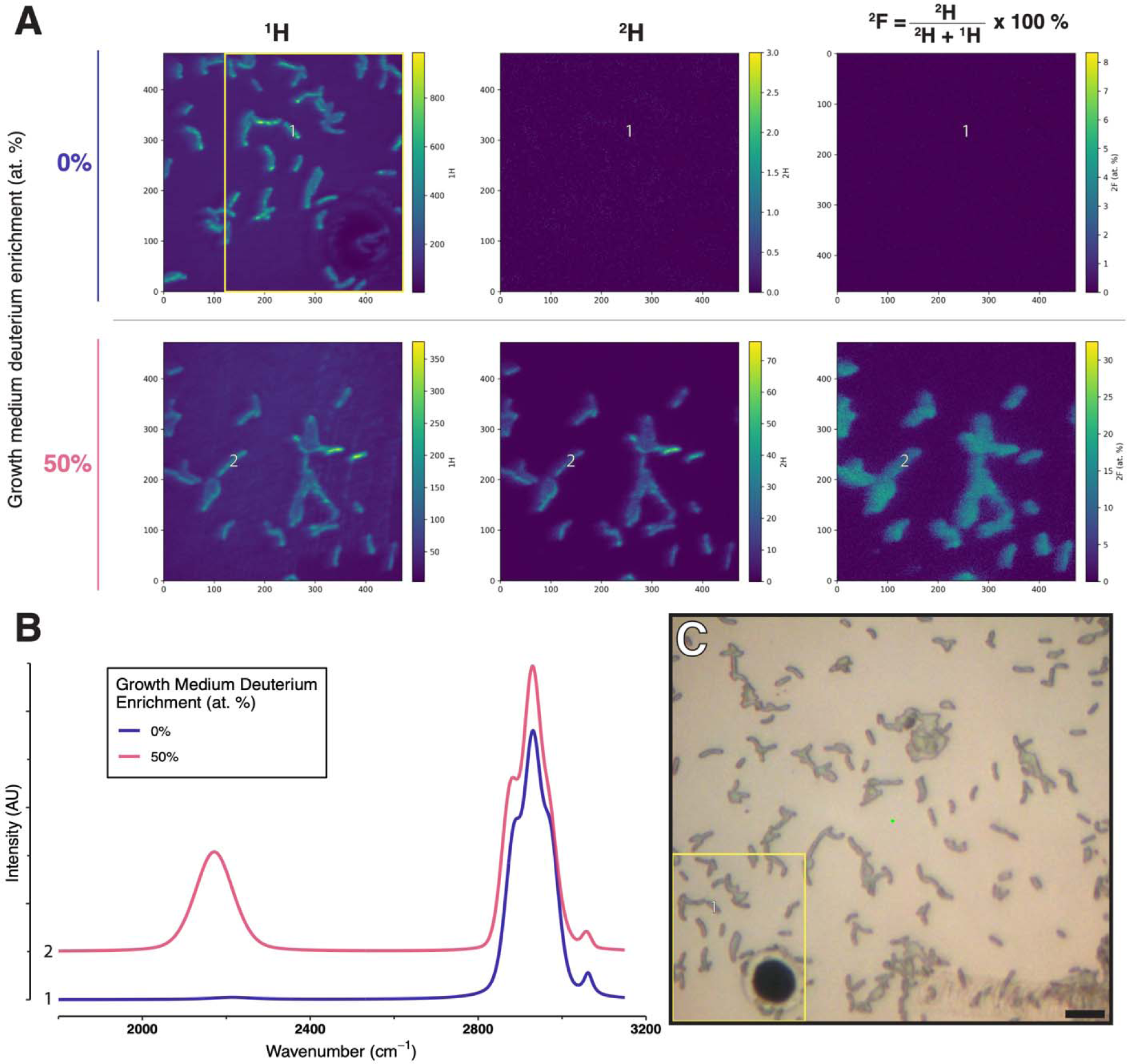
Representative Raman and nanoSIMS data collected in this study. Across all panels data from two representative cells, “1” and “2”, are noted. Representative nanoSIMS isotopic images of are shown in **Panel A**, with pixel counts labeled on the x and y axes. From left to right, the pixel intensity in each panel corresponds to direct ion counts for ^1^H^-^, ^2^H^-^, and the fractional abundanc of deuterium (*^2^F*), respectively. The top row shows cells grown in 0% ^2^H_2_O (natural abundance); th bottom row displays cells grown in 50 at. % ^2^H_2_O. Representative single-cell fitted Raman spectra are shown in **Panel B**, with the characteristic C-H band between 2800 and 3000 cm^-1^ clearly visible. For cells grown in deuterated media (Cell 2), the C-D band emerges between 2040 and 2300 cm^-1^. **Panel C** displays a typical micrograph captured through a confocal Raman microspectroscope. The scale bar represents 2µm. Cells in all panels represent *T. hydrogeniphilus.* The yellow boxes in the top-left facet of Panel A, and Panel C, represent a correlated region imaged with nanoSIMS and Raman (reflected light), respectively.

For our study, we built upon this prior work in four key aspects. First, we generated a robust hydrogen isotopic calibration for Raman spectroscopy that specifically targets two environmentally-relevant taxa (**Fig. 2**). Second, we combined the results of our isotopic calibration with a numerical model of microbial growth to determine the feasibility and sensitivity of Raman-SIP for quantitative inference of microbial biomass growth rate (**Fig. 3**). The ability to translate Raman-derived measurements of deuterium enrichment into biomass synthesis/growth estimates would pave the way for future investigations to not only identify microbial activity *in situ*, but also to quantify biosynthetic growth rate. Third, the cells used to generate our Raman-based isotopic calibration were correlatively measured by nanoSIMS to determine the extent to which nanoSIMS and Raman-based measurements agree. Finally, we provide a computational framework for Raman-SIP that allows users to optimize SIP incubations for their specific study system.

**Figure 2:**
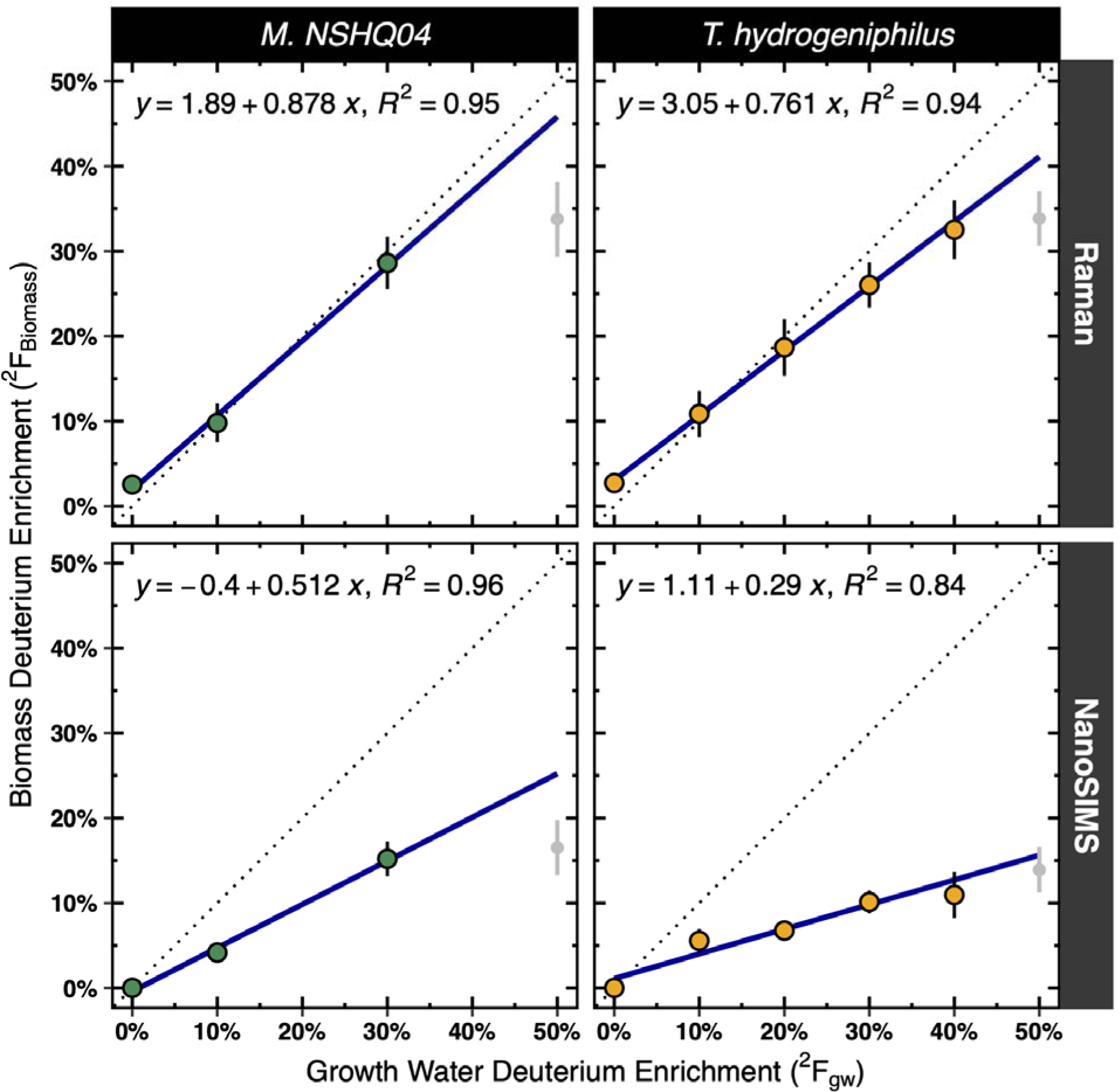
Hydrogen isotope calibrations relating growth water and biomass ^2^H content. Individual measurements of single-cell biomass deuterium enrichment (*^2^F_biomass_*) conducted with Raman spectroscopy (top panels) or nanoSIMS (bottom panels) are plotted against the corresponding deuterium content of the microbial growth media (*^2^F_gw_*). The dotted line indicates the 1:1 line; the solid blue lin represents the line of best fit whose equation is noted in each panel. Points and error bars displayed are mean and standard deviation of biomass deuterium enrichment, respectively. Note that the linear models displayed in each panel are calculated only with data between deuterium enrichments of 0 – 40 at. % to ensure that these models are not affected by the toxicity effects observed at 50 at. % label isotopic enrichment (point intervals in gray).

**Figure 3:**
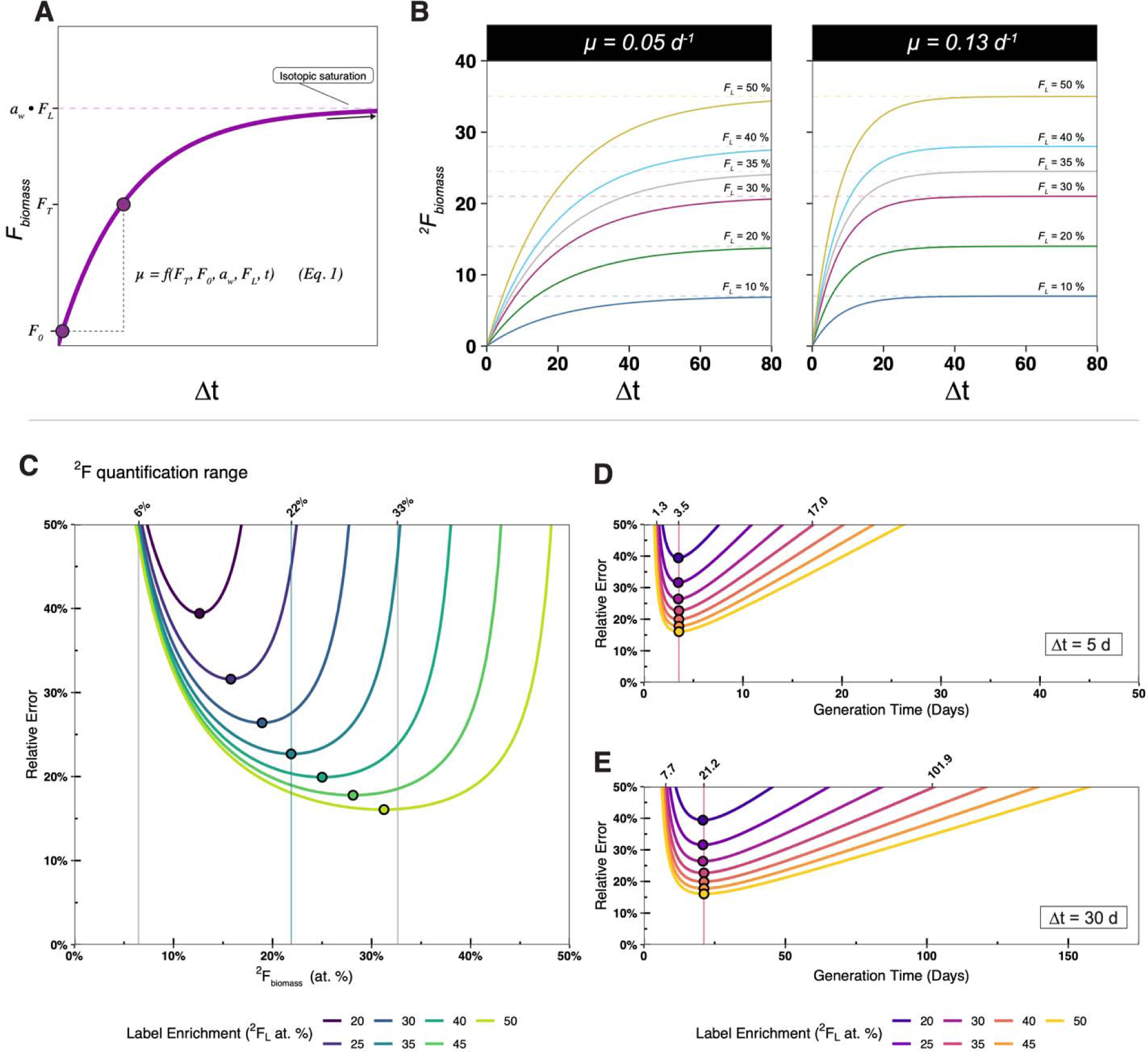
Modeling of microbial growth in the presence of enriched isotopic label (A, B) and associated error in biomass ^2^F quantification (C) and growth rate/generation time (D, E). A schematic figure (**Panel A**) visualizes how microbial growth rate is calculated as a function of biomass isotopic enrichment (*F_T_*), label strength (*F_L_*), incubation time (*t*), and water assimilation efficiency constant (*a_w_*) (**Eq. 1**). Similarly, **Panel B** displays example output of the microbial growth model, where *a_w_* = 0.7. Isotopic enrichment increases as a function of time but depends upon isotopic label strength (*F_L_*) and growth rate (*µ*, noted above each plot). In **Panels C-E**, we display quantification ranges where relative error in the growth rate measurement is < 50% of the growth rate itself (Relative Error = σ*_µ_ / µ × 100%* where µ is apparent growth rate). In **Panel C,** relative error is plotted against biomass deuterium enrichment (*^2^F_biomass_*) for label strengths *^2^F_L_*ranging from 20-50 at. %. Plotted points indicate the of minimum error for each isotopic label. Along the top axis of **Panel C**, the lower limit, error optimum, and upper limit of quantification, are noted, for the isotopic label strength of *^2^F_L_*= 35 at. %. The key results from **Panel C** are summarized in **Table 2**. In **Panels D** and **E,** relative error is plotted against biomass generation times in the context of a simulated SIP incubation. In these examples, the incubation times of 5 days (**Panel D**) and 30 days (**Panel E**) are chosen. Along the top axes of **Panels D** and **E**, the minimum, optimum, and maximum quantifiable generation times are noted for the isotopic tracer strength of *^2^F_L_* = 35 at. %. These generation times correspond to the limits of quantification and optima noted in Panel A.

## Results and Discussion

### Single-cell isotopic measurements

To assess the sensitivity and precision of Raman microspectroscopy as a reporter of cellular ^2^H incorporation, we generated a large isotopic dataset of bacterial and archaeal cells grown to equilibrium with deuterated (^2^H_2_O) media of varying hydrogen isotope compositions (0 – 50 at. % ^2^H). We focused our study on two anaerobic organisms isolated from the subsurface biosphere: a sulfate reducing bacterium (SRB) *Thermodesulfovibrio hydrogeniphilus* (43), and a methanogenic archaeon *Methanobacterium NSHQ04* (44). We acquired individual measurements of cellular deuterium enrichment (n = 351) with Raman spectroscopy and nanoSIMS (**Fig. 1**). We refer to hydrogen isotope content using both Raman and nanoSIMS as the fractional abundance of deuterium, *^2^F*, reported in atom percent (at. %) (see *Materials and Methods*), where *^2^F =* ^2^H / (H + ^2^H) *× 100 %.* To generate hydrogen isotope calibrations, we fit simple linear models relating the hydrogen isotopic composition of microbial biomass (*^2^F_biomass_*) to the organism’s growth water (*^2^F_gw_*). (**Fig. 2**).

With Raman microspectroscopy, biomass ^2^H enrichment (*^2^F_biomass_*) increased linearly and exhibited strong positive correlation (R^2^ = 0.94, 0.95, for *T. hydrogeniphilus and M. NSHQ04,* respectively) with the deuterium content of the growth water (*^2^F_gw_*) (**Fig. 2**, **Table 1B**). Raman measurements of biomass deuterium content exhibited an average standard deviation of σ_2F_ = 2.89 at. %. The variability of Raman-derived *^2^F_biomass_* increased with the isotopic composition of the growth water: a maximum standard deviation was observed at the *^2^F_gw_* = 50% ^2^H_2_O condition where σ_2F_ = 3.77 at. %. Corresponding measurements by nanoSIMS of the same cells followed a similar trend: *^2^F_Biomass_* increased linearly with the growth water applied (R^2^ = 0.84, 0.96 for *T. hydrogeniphilus* and *M. NSHQ04*, respectively) and the average standard deviation of nanoSIMS *^2^F* was σ_2F_ = 1.86 at. %. Similar to Raman, the standard deviation of nanoSIMS measurements increased along with the isotopic composition of growth water, reaching a maximum at the 50% label where σ_2F_ = 3.20 at. % (**Table 1A**). With both analytical techniques and across both organisms, we observed depressed biomass enrichments at *^2^F_gw_* = 50 at. %, likely resulting from deuterium toxicity effects (see section *Applying Raman microspectroscopy to SIP growth rate measurements*). For analytical uncertainty, we report the average standard deviation observed in Raman and nanoSIMS measurements across growth water conditions: 2.89 and 1.86 at. %, respectively. We emphasize that these are conservative over-estimates of analytical error due to heterogeneity in the hydrogen isotopic enrichment of individual cells. Our data support previous observations by Berry et al. that point to high reproducibility of single-cell measurements (8).

**Table 1.** Results and statistical summary of the isotopic calibration. Here we summarize key outputs of our analyses: (A) analytical uncertainty in biomass measurements across growth water isotopic compositions (^2^F_gw_), (B) Linear models relating the isotopic composition of biomass (^2^F_biomass_) to that of growth water (^2^F_gw_), (C) Water hydrogen assimilation efficiency factors (*a_w_*) derived from the hydrogen isotopic calibration, with associated standard error, (D) Assessment of correlation between biomass isotopic values estimated with Raman spectroscopy (*^2^F_Raman_*)and nanoSIMS (*^2^F_nanoSIMS_*) (Supplementary Fig. S2, S3)

|  | Raman | nanoSIMS |
| --- | --- | --- |
| **(A) Analytical Uncertainty in Cellular $^2F$ (SD) (at. %)** |  |  |
| $^2F_{gw} = 0\%$ (SD of the method blank) | 1.13 | 0.01 |
| $^2F_{gw} = 10\%$ | 2.56 | 1.24 |
| $^2F_{gw} = 20\%$ | 3.34 | 1.04 |
| $^2F_{gw} = 30\%$ | 3.07 | 2.95 |
| $^2F_{gw} = 40\%$ | 3.46 | 2.74 |
| $^2F_{gw} = 50\%$ | 3.78 | 3.20 |
| Mean SD across all conditions | 2.89 | 1.86 |
| **(B) Hydrogen Isotope Calibration Models** |  |  |
| where: $y = ^2F_{Biomass}$, $x = ^2F_{gw}$ | | |
| *T. hydrogeniphilus* | $y = 0.761x + 3.05$<br>$R^2 = 0.94$ | $y = 0.29x + 1.11$<br>$R^2 = 0.84$ |
| *M. NSHQ04* | $y = 0.878x + 1.89$<br>$R^2 = 0.95$ | $y = 0.512x - 0.40$<br>$R^2 = 0.96$ |
| **(C) Water hydrogen assimilation efficiency** |  |  |
| $a_w \pm SE$ | | |
| *T. hydrogeniphilus* | $0.761 \pm 0.014$ | $0.290 \pm 0.009$ |
| *M. NSHQ04* | $0.878 \pm 0.020$ | $0.512 \pm 0.010$ |
| **(D) Correlative Statistics** |  |  |
| Linear model equation<br>*where:*<br>$y = {}^2F_{nanoSIMS}$<br>$x = {}^2F_{Raman}$<br>(Supplementary Text, Supplementary Fig. S2-S3) | $y = 0.439x - 0.3617$<br>Slope SE = 0.01<br>Intercept SE = 0.24 | |
| RMSE (at. %) | 2.52 |  |

### Applying Raman microspectroscopy to SIP growth rate measurements

Having generated a Raman-based hydrogen isotopic calibration of microbial biomass, we sought to test the applicability of this analytical method to measurements of cell-specific growth rate. Microbial growth rate can be inferred from SIP incubations because the degree of an organism’s biomass isotopic enrichment over time depends on the isotopic composition of a given substrate and its rate of biomass synthesis (14). Therefore, if measurements of ^2^H incorporation are reliable, then these values should translate to cell-specific growth rates. To this end, we sought to parameterize sources of uncertainty and propagation to downstream calculations to define acceptable experimental conditions and analytical uncertainties for quantitative measurement of microbial growth rate.

The growth rate of an organism in the presence of a stable isotope tracer is calculated with **Eq. 1** (14). In brief, microbial growth rate is a logarithmic relationship between an organism’s isotopic composition at the start of an incubation (*F_0_*) and its isotopic composition at the time (*t*) of sampling (*F_T_*). Cellular isotopic enrichment is compared to the enrichment of the isotopic label (*F_L_*) offset by a water hydrogen assimilation efficiency constant (*a_w_*) – a value representing the fraction of biomass hydrogen that a cell acquires from its growth water, as opposed to other metabolic sources and associated fractionation effects.

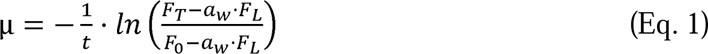

We modeled growth rates (*µ*) (**Eq. 1**) across a wide range of SIP experimental conditions (varying *F_L_*, *t,* and *a_w_*,) and applied standard propagation of uncertainty to determine the associated errors (σ_µ_) (**Eq. 2**) that result from compounding uncertainty of Raman-SIP experimental aspects (3,14):

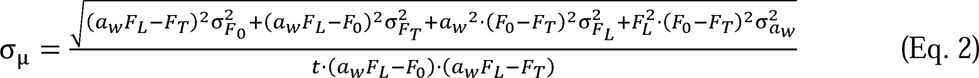

Where sigma (σ) terms represent uncertainties of each subscripted parameter (e.g., σ*_FT_* represents uncertainty in biomass isotopic composition at time of sampling *T*). We convert growth rate to biomass generation time using the relationship:

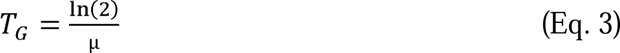

where *µ* is growth rate in units of *t^-1^* and T_G_ is generation time in units of *t*.

To a first order, the range of growth rates (*µ*) that can be measured with SIP depends on the isotopic composition of the label and the incubation duration. Uncertainty in growth rate (σ*_µ_*) depends on analytical uncertainty in label strength (σ*_FL_*), analytical uncertainty of biomass isotopic enrichment (σ*_FT_*), and microbial water hydrogen assimilation efficiency (σ*_aw_*). Our model constrains the region of quantification for both isotopic enrichment (**Fig. 3C**) and corresponding growth rate (**Fig. 3D-E**) in Raman SIP experiments. We propagate the uncertainty estimates of ^2^*F_biomass_* estimated from our hydrogen isotope calibration as σ*_FT_* to estimate associated uncertainty in growth rate (σ_µ_) that results from a given SIP experiment. We report uncertainty in growth rate relative to the growth rate itself as *Relative Error = (*σ*_µ_ / µ) × 100%*. We set our limit for relative error to be 50% such that errors below this cutoff result in growth rate differences that are distinguishable with > 2σ confidence.

In **Fig. 3** we display example outputs from our model using incubation times of *t = 5* and *t = 30 days* across a range of ^2^H_2_O label strengths. For example, the range of quantification for an experiment applying a 35% label is 6.9 - 32.6 at. % deuterium enrichment. Ranges of quantification and error optima for additional isotopic label strengths are reported in **Table 2**. For a 5-day incubation, this range of *^2^F* quantification corresponds to growth rates between 0.53 – 0.04 *d^-1^,* or generation times of 1.3 and 17 days. For a 30-day incubation, this range of *^2^F* quantification corresponds to growth rates between 0.09 - 0.007 *d^-1^* or generation times between 7.7 and 101.9 days (**Fig. 3E**).

**Table 2:** Range of Quantification for Raman-derived (CD%-based) measurements of single-cell deuterium content (**Fig. 3C**). The optimum value represents the ^2^F_biomass_ value at which relative error in growth rate calculation is minimized. The lower and upper limits of quantification represent where relative error of growth rate calculation exceeds 50%.

| Isotopic label strength ($^2F_L$ at. %) | Lower limit of quantification ($^2F_{\text{biomass}}$ at. %) | Optimum ($^2F_{\text{biomass}}$ at. %) | Upper limit of quantification ($^2F_{\text{biomass}}$ at. %) |
| --- | --- | --- | --- |
| 20 | 7.4 | 12.6 | 16.7 |
| 25 | 6.8 | 15.8 | 22.4 |
| 30 | 6.6 | 18.9 | 27.7 |
| 35 | 6.4 | 21.8 | 32.6 |
| 40 | 6.4 | 25.0 | 37.7 |
| 45 | 6.3 | 28.1 | 42.8 |
| 50 | 6.2 | 31.3 | 48.0 |

Asymptotic rises in uncertainty at high and low biomass isotopic enrichment, respectively, correspond to (i) uncertainty in growth rate as cell isotopic enrichment approaches that of the label solution and (ii) uncertainty resulting from biomass isotopic signal not significantly exceeding that of the noise inherent to the analytical method. The first condition (i) represents cells that grow faster than can be discriminated using the set incubation time and label strength, while (ii) represents cells that grow too slow to be distinguished from noise. It should be noted that our ranges of quantification should be viewed conservative estimates, as the source of uncertainty we apply corresponds to heterogeneity of ^2^H-incorporation across homogenous cell populations, as opposed to the error across individual Raman acquisitions. This latter source of error is far smaller both in our dataset and in previously reported datasets (8).

Incubation time (*t*) shifts the dynamic range of SIP (**Supplementary Fig. S8**): shorter incubations can resolve faster-growing organisms from each other but may not capture slower-growing organisms due to insufficient label incorporation. Conversely, longer incubations can capture slower-growing organisms, but faster-growing organisms may become saturated with the label and will no longer be resolvable from each other (**Fig. 3**). Growth rates of fast-growing organisms, those that are readily saturated with deuterium, could lead to underestimates of aggregate microbial turnover.

Increasing the strength of the label (*F_L_*) expands the dynamic range of the SIP method i.e., both slower and faster growth rates can be captured as there is more isotopic “space” that can be measured (**Table 2**, **Fig. 3**). However, increasing *F_L_* must be weighed against the risk that higher deuterium concentrations can impose toxicity or unintended physiological effects. Deuterated water at high levels of isotopic labeling have been shown to have no effect, inhibition and even stimulation among different microorganisms (8,10,45). In our experiments, we observed depressed isotopic enrichment in growth water containing *50 at. %* ^2^H_2_O. These effects were noted for both organisms and both the Raman and nanoSIMS data (grey point intervals in **Fig. 2**). We conjecture that this result stems from isotopic toxicity affecting microbial growth or metabolic switching at high isotopic enrichment, but this mechanism cannot be explained by our study. Therefore, we caution against applying ^2^H_2_O label strengths at or above 50 at. % due to varying physiological effects that may arise at this threshold, especially when applying this approach to natural microbial communities of unknown and mixed composition. Uncertainties in isotopic label strength (σ*_FL_*), especially at high isotopic enrichment, due to pipetting or gravimetric dilution contribute relatively minor components to total uncertainty.

The water hydrogen assimilation efficiency constant (*a_w_*) is essential for estimation of microbial growth rate, as it sets the upper limit of biomass deuterium enrichment. This limit is represented by the *a_w_ F_L_* terms in **Eq. 1** (**Fig 3A**). Therefore, similar to increasing the strength of the label, an organism with a greater *a_w_* has more isotopic “space” in which to become enriched. In our study, we use Raman-derived *^2^F* measurements to empirically determine the *a_w_* of *T. hydrogeniphilus* and *M. NSHQ04* to be *a_w_ = 0.*76 ± 0.02 and *a_w_ = 0.88* ± 0.02, respectively (**Table 1C**, **Fig. 2**). The higher *a_w_* observed with the methanogen *M. NSHQ04* is in line with primarily autotrophic growth (44) with an offset from *a_w_* = 1 due to hydrogen isotopic fractionation. The slightly lower *a_w_* for the sulfate reducing *T. hydrogeniphilus* is indicative of a hydrogen contribution from its acetotrophic metabolism or from greater fractionation effects. We note that these values represent *a_w_* of whole-cell biomass, which may be distinct from those calculated via compound-specific analysis (3,21,25). It was previously noted that uncertainty in *a_w_* is a significant driver of uncertainty in hydrogen SIP-derived growth rate estimates (3). For mixed communities, this uncertainty must be propagated through growth rate calculations. For organisms with known metabolisms, this term can be estimated as common metabolic modes often constrain *a_w_* factor to a degree (3,21,25).

### Raman and nanoSIMS measure different pools of cellular hydrogen

Currently, nanoscale secondary ion mass spectrometry (nanoSIMS) serves as a standard method for reporting single-cell isotope ratios. A NanoSIMS instrument applies a primary ion beam to ionize and ablate a sample, inducing the release of secondary ions whose isotopic composition is measured by magnetic sector isotope ratio mass spectrometry. NanoSIMS has many benefits as an analytical technique: it can measure multiple isotopes simultaneously and allows spatial/elemental imaging at the nanoscale. However, nanoSIMS destroys the sample and is time and cost-intensive. Furthermore, depression of isotopic enrichment measured by nanoSIMS due to sample preparation has been widely reported for multiple stable isotope tracers (^13^C, ^18^O, ^15^N, ^2^H, and ^34^S) under various fixation and staining conditions (8,10,15,36,40–42,46). Given these limitations, it would be useful to have additional methods that can complement the strengths of nanoSIMS while negating some of its drawbacks.

Deuterium enrichments measured by Raman were greater than those measured by nanoSIMS, when compared to the culture growth water (**Fig. 2**). When we correlated cell-specific *^2^F_biomass_* measurements we observed that, cell-to-cell, nanoSIMS-derived hydrogen isotope values were severely depressed compared to corresponding Raman-derived values (**Supplemental Fig. 2**). From this correlative dataset, we calculated an average hydrogen isotope dilution factor of 58.7 ± 6.0 % across all growth water labeling conditions (*Supplementary Text*).

Such a significant dilution of hydrogen isotopes speaks to a fundamental difference in the nature of nanoSIMS- and Raman-based measurements. Owing to how the primary ion beam ionizes and ablates a cell, nanoSIMS is mostly agnostic to the sources of ^2^H/^1^H it measures, setting aside potential differences in hydrogen ionization efficiency across different classes of molecules. Therefore, ^1^H^-^ and ^2^H^-^ ions reaching the instrument detectors can derive from a variety of cellular hydrogen sources including remnant intracellular water, adsorbed extracellular water, and biomass hydrogen in both non-exchangeable sites and exchangeable sites that are readily mixed with natural-abundance washing buffers during sample preparation. Protonated sites in biomolecules (e.g., O-H, N-H, S-H, etc.) experience rapid exchange with aqueous wash solutions on the order of picoseconds to minutes (47,48), meaning that anabolically produced O-^2^H, N-^2^H, S-^2^H, etc. bonds would be rapidly overprinted by natural-abundance hydrogen during standard washing procedures (**Fig. 4**). As described in *Materials and Methods*, no stains or probes were applied to the cells in this study. Therefore, the hydrogen dilution factors reported here result solely from sample washing. This significant and persistent isotopic dilution may confound efforts to use deuterium tracers as measures of biosynthesis in nanoSIMS-based studies (10). This problem is compounded at lower-isotopic enrichment, where substantial variability in ^2^H/^1^H values results from low ^2^H ion counts as well as random variation in which cell subcomponents (e.g., highly exchangeable vs. non-exchangeable) are ablated (10). Our observation that the magnitude of isotopic dilution appears variable across samples (**Fig. 2**) suggests that dilution factors could contribute substantial uncertainty to corrected values. Future work should apply caution when applying dilution factors to back-calculate isotopic abundance, as these can vary widely between the sample preparation and organism (40), an effect that may be pronounced in environmental systems with mixed microbial communities.

**Figure 4.**
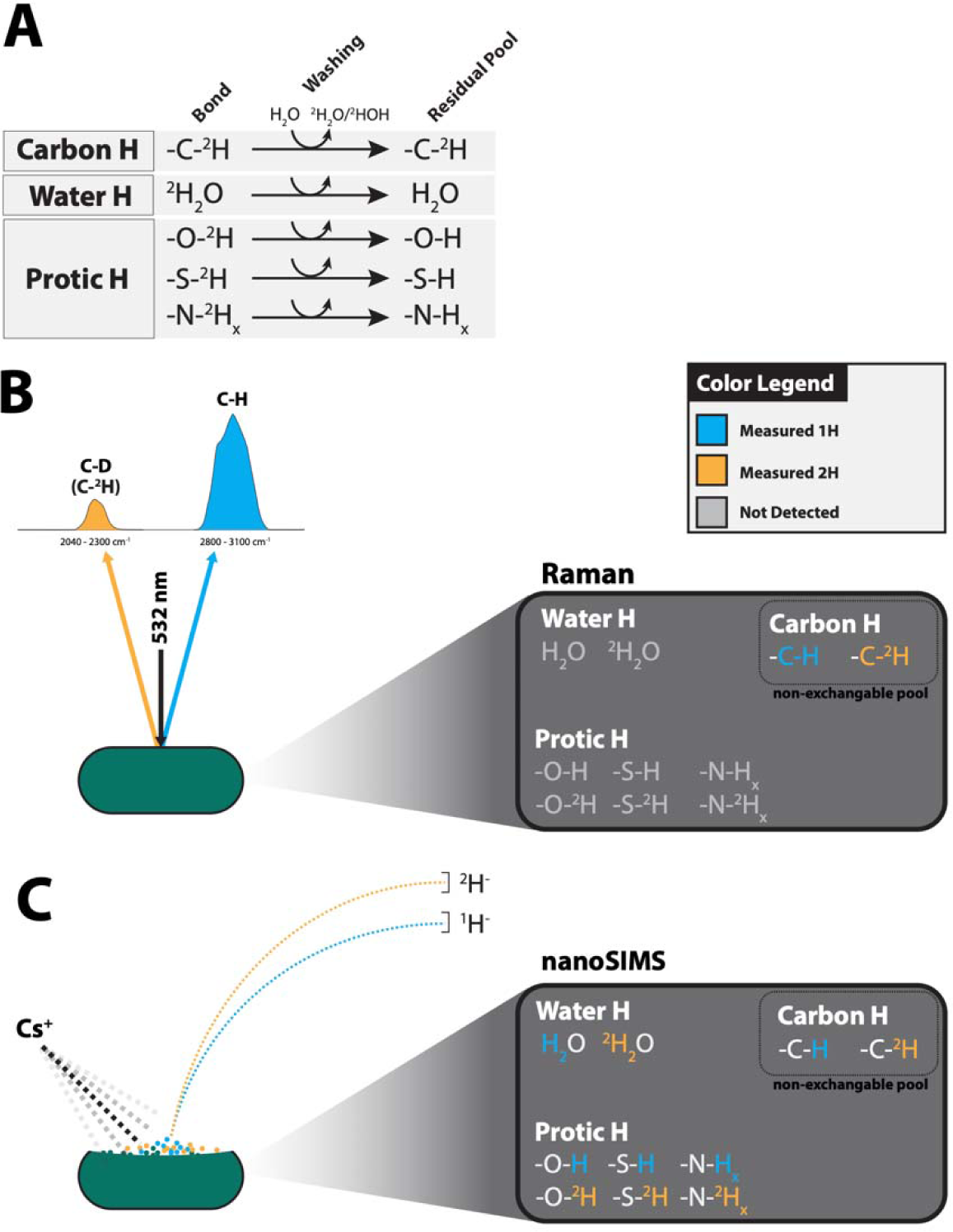
Schematic diagram showing the dilution of ^2^H/H isotopic signal due to sample preparation, and how Raman and nanoSIMS measure different pools of biomass ^2^H/H. (A) The original pool of deuterated cell mass is biosynthesized, sourcing ^2^H from the growth medium. During sample washing, hydrogen in exchangeable, or “protic” bonding environments rapidly exchange with wash solutions, leaving a diluted residual pool of ^2^H/H. Carbon-hydrogen and carbon-deuterium bonds are not affected by this dilution effect, as ^2^H/H in these bonds do not exchange during washing. **(B)** Within the C-H and C-D (C-^2^H) wavenumber regions, Raman is only sensitive to carbon-hydrogen and carbon-deuterium bonds, which are not affected by dilution due to washing, whereas **(C)** nanoSIMS ostensibly measures all pools of ^2^H/H, including those affected by dilution.

A key advantage of Raman-SIP is that the C-H (2800 - 3100 cm^-1^) and C-D (2040 - 2300 cm^-1^) bands report solely non-exchangeable carbon-hydrogen and carbon-deuterium stretching modes associated with mostly lipids, proteins and nucleic acids (8,49). These spectral regions are oblivious to exchangeable bonding environments including O-H, N-H, S-H, etc., as well as remnant water. Raman spectroscopy therefore focuses on biosynthetically produced organic bonds without the confounding effects of remnant water or exchanged biomolecular hydrogen. Deuterium measurements by Raman-SIP appear to exhibit a greater reliability in tracking biosynthetic incorporation of deuterium into cellular biomass.

### Future Directions for Raman-SIP

Raman spectroscopy has several advantages for measuring microbial growth rates. First, it is a non-destructive, rapid method that is readily coupled with other techniques such as fluorescence microscopy, fluorescence in situ hybridization (FISH), scanning electron microscopy (SEM), or cell sorting (8,15). A single Raman spectrum takes between seconds to one minute to acquire, and many Raman instruments are equipped with fluorescence modules that can enable visualization of the cells in addition to transmitted or reflected light microscopy. Second, Raman is comparatively accessible relative to nanoSIMS and allows analysis with minimal preparation, making it capable of being deployed directly on many forms of microbial systems including filters from aquatic environments, rock faces, plant material, etc. Third, Raman microspectroscopy has microscale spatial resolution and provides diagnostic information about cellular organic composition. The fingerprint region (200 – 1800 cm^-1^) is routinely used to identify biomolecules (49) as well as differentiate taxa, down to the species/strain level and growth stage (50–52). Fourth, the Raman fingerprint region enables identification of biologically precipitated minerals and the mineralogical/material context of a cell (37,53). Raman SIP with deuterium, specifically, makes identification of individual cell and mineral components in organic-mineral assemblages feasible because the C-D band occupies a typically silent region in inorganic Raman spectra. Co-registered mineral and quantitative cell-activity measurements open the door for novel investigations into the cell-mineral and cell-cell relationships that drive microbial activity in the rock-hosted biosphere (35). Furthermore, measurement of ^13^C and ^15^N enrichment in microbial cells has been demonstrated with Raman (55–59) and the potential exists for the co-registration of multiple stable isotopes at the single-cell level.

The main disadvantage of deuterium Raman-SIP is its detection limit. Classical or spontaneous Raman microspectroscopy of microbial cells is limited by low signal intensities, particularly in samples with high autofluorescence (54). A key outcome of our study is the definition of Raman-SIP ranges of quantification. Our study defines the minimum cutoff for quantitative measurement as ∼ 7 at. % (**Table 2**), depending on the label strength applied. In the coming years, advancements in Raman technology hold potential for increasing its sensitivity and subsequently improving isotopic measurements. Specifically, surface- or tip-enhanced Raman spectroscopy (SERS or TERS) (50,60–63), stimulated Raman scattering (SRS), resonance Raman spectroscopy, or coherent anti-Stokes Raman spectroscopy (CARS), which are shown to dramatically improve acquisition times, and enhance signal, could be applied to determine isotopic enrichments (55,56,60,64–69). However, at present, robust validation of these methods for deuterium SIP is required (**Supplemental Text, Supplemental Fig. S4 – S5**).

Our results point to Raman microspectroscopy being a powerful tool for conducting SIP, especially where complex environmental matrices may complicate bulk-omics or other single-cell approaches. For the benefit of future researchers, we designed an interactive graphical user interface (GUI), *Shiny R-SIP*, that allows users to implement our SIP model and optimize their experiments by considering how the relevant sources of uncertainty impact the design of a Raman-SIP experiment (*Supplementary Data* or online: https://apps.kopflab.org/login with login credentials username: *shiny-rSIP*, password: *public-access*). The GUI allows users to adjust parameters of label strength, incubation time, and water hydrogen assimilation efficiency, as well associated uncertainties in these terms, in order to design effective SIP experiments. The parameters we define in our microbial growth model are specific to the study organisms used and our Raman instrument: they are intended to provide reasonable estimates of uncertainty that may be encountered. Users of this GUI are encouraged to define their own uncertainty terms by constraining the technical variation in individual CD% measurements inherent to their Raman instrument. For samples where microbial growth may exhibit large variation, we suggest users consider multiple incubation times (or sub-sampling efforts) to effectively capture a wide range of biomass growth rates in experiments targeting natural microbial communities, while also quantifying their threshold of tracer saturation as an upper-limit of growth-rate quantification.

### Conclusions

In this work, we provide the basis for the quantitative measurement of single-cell microbial growth rates via Raman-SIP. We develop hydrogen isotope calibrations for two organisms, *T. hydrogeniphilus* (sulfate-reducing bacterium) and *M. NSHQ04* (methanogenic archaeon). These calibrations validate the utility of Raman microspectroscopy as a method for measuring microbial ^2^H/H enrichment. Applying our Raman-derived isotopic calibration to a model of microbial growth, we define ranges of quantification for Raman SIP experiments where single-cell growth rates can be sensitively distinguished. We find that with reasonable (20 - 40 at. % ^2^H_2_O) isotopic label strengths and incubation times, Raman-SIP can capture a wide array of microbial generation times, with this range being defined by the parameters of the SIP experiment. These ranges of quantification can guide the design and interpretation of SIP experiments where Raman is used to track cellular ^2^H incorporation. Finally, we observe that hydrogen isotopic values derived from nanoSIMS suffer from severe dilution that may complicate the use of nanoSIMS for quantifying cellular ^2^H uptake. In conclusion, this work provides a robust framework for applying deuterium Raman-SIP to spatially resolved, quantitative investigations of microbial activity in environmental and model systems.

## Materials and Methods

### Microorganisms and Growth Conditions

We cultivated *Thermodesulfovibrio hydrogeniphilus* (43) and *Methanobacterium NSHQ04* (44), in media containing 0%, 10%, 20%, 30%, 40%, or 50% ^2^H_2_O (0% = local ultra-pure water which contains ∼140ppm ^2^H_2_O naturally). Base media (described below) were modified with different percentages (vol/vol) of ^2^H_2_O [Cambridge Isotope Laboratories], depending on the final isotopic enrichment desired for the experiment. All cultures were grown in the dark in anaerobic 60 ml serum vials containing 25 ml of growth medium.

*Thermodesulfovibrio hydrogeniphilus* HBr5^T^ (DSM 18151) (43), a sulfate reducing bacterium, was obtained from the Deutsche Sammlung von Mikroorganismen und Zellkulturen (DSMZ). It was grown in a modified DSM 641 medium that contained (per liter): 1.0g NH_4_Cl, 2.0g Na_2_SO_4_, 1.0g Na_2_S_2_O_3_x5H_2_O, 1.0g MgSO_4_x7H_2_O, 0.1g CaCl_2_x2H_2_O, 0.5g KH_2_PO_4_, 2.0g NaHCO_3_, 1ml trace element solution SL-10, 1ml selenite-tungstate solution, 1g yeast extract, 0.5ml sodium resazurin (0.1% wt/vol), 0.2g sodium acetate, 10ml 141 vitamin solution, and 0.1g Na_2_S x 9H_2_O. The headspace was flushed and over-pressurized with 1 bar H_2_:CO_2_ (80:20).

*Methanobacterium NSHQ04* (44) is a methanogenic archaeon enriched from groundwater isolated from the Samail ophiolite in Oman. The culture was grown in a synthetic site water (NSHQ04) medium (44) containing (per liter): 420mg NaCl, 683mg CaCl_2_x2H_2_O, 1.2mg H_4_SiO_4_, 1.7mg NaBr, 60mg MgCl_2_x6H_2_O, 100mg NH_4_Cl, 10 ml 141 trace elements, 10ml 141 vitamins, 0.5ml sodium resazurin (0.1 wt/vol), 20ml antibiotic cocktail [containing per 100ml MilliQ water, 500mg penicillin G, 500mg streptomycin, 500ul ampicillin (10mg/ml stock)], 10ml KH_2_PO_4_ (1.1 wt/vol), 4ml yeast extract (5% wt/vol), 2ml Fe(NH_4_)_2_(SO_4_)_2_ (0.2% w/v), 10ml sodium formate (100mM), 10ml cysteine-HCl (2.5% w/v), and 10ml Na_2_S (2.5% w/v). The headspace was flushed and over-pressurized with 2 bar H_2_:N_2_ (5:95).

Cultures of *T. hydrogeniphilus* and *M. NSHQ04* were incubated in forced-air incubators at 65°C and 40°C, respectively. Cultures were transferred three times into media of identical deuterium enrichment. Before each transfer, cultures were grown to stationary phase, which was reached in 3-4 days for *T. hydrogeniphilus* and 12-14 days for *Methanobacterium NSHQ04*. This ensured that the cells measured were at or near isotopic equilibrium with their respective growth water, offset by *a_w_*. After the third transfer, 1ml of each culture was harvested and fixed by addition of paraformaldehyde to a final concentration of 2% (vol/vol). Fixed cells were washed by centrifugation in successively dilute phosphate-buffered saline (PBS) of rinses [1X, 1%, 0.1%, and 0.01% (vol/vol) PBS) to ensure the removal of excess salts and fixative. PBS was composed of Dulbecco’s formula: KCl 200 mg/L, KH_2_PO_4_ 200 mg/L, NaCl 8000 mg/L, Na_2_HPO_4_ 1150 mg/L. After the final wash, cells were resuspended in 50μl of 0.01% (vol/vol) PBS.

### Sample Preparation

Sample coupons were prepared by raster engraving of aluminum-coated glass slides (Deposition Research Lab Inc.) with an Epilog Mini 24 CO_2_ laser machine operating at 80 W laser power, 10% etching speed, to create a 3 x 6 grid. These etchings were used to manually break the glass slide, producing 7mm^2^ coupons that are compatible with both Raman and nanoSIMS sample mounts. Aluminum coupons were used instead of silicon wafers (typically used for nanoSIMS studies) because silicon exhibits a large peak at 520 cm^-1^ in the Raman spectrum. All coupons were sterilized and rendered organic-clean by combustion for 8 hours at 450°C in a muffle furnace. 5μl of fixed and washed cells was spotted on individual coupons and allowed to air-dry.

### Raman Spectroscopy, Fitting, and SIP Calculations

Raman spectroscopy was conducted at the Raman Microspectroscopy Laboratory, Department of Geological Sciences, University of Colorado-Boulder (RRID:SCR_019305) on a Horiba LabRAM HR Evolution Raman spectrometer equipped with a 100mW 532 nm excitation laser. The laser was focused with a 100X (NA = 0.90) air objective lens, resulting in a spot size of ∼1µm. Single-cell spectra were captured in the 200-3150 cm^-1^ range using 100% laser power (2.55 mW) over 2 acquisitions of 45 seconds. A 100 µm confocal pinhole and 600 lines/mm diffraction grating were used, resulting in a spectral resolution of ∼ 4.5 cm^-1^. Duplicate acquisitions were averaged to remove noise and cosmic ray spikes. Spectra were baseline-subtracted using a polynomial fit in LabSpec 6 (Horiba Scientific). Fitting of bands in the 1800 - 3150 cm^-1^ region was performed using Gaussian-Lorentzian curve fitting algorithms to integrate C-D (2040 – 2300 cm^-1^) and C-H (2800 – 3100 cm^-1^) bands. Raman-derived *^2^F* was determined by calculating the CD% value of each spectrum: *^2^F = CD% = CD / (CH + CD) × 100%* where CD and CH are the areas of the C-D and C-H bands, respectively.

To correlate Raman spectra with nanoSIMS data, extensive context maps of the sample coupons were acquired using reflected light microscopy on the Raman instrument. Before cell spotting, fiducial markings were etched into the aluminum coupon surface with a Leica LMD7000 microdissection system.

As described in *Results and Discussion*, modeling of growth rate was carried out with **Eq. 1–3**. (3,14). Using these formulae, we modeled *µ* across a range of *F_L_*, *t*, and *a_w_*. We define an acceptable cutoff using a relative error term defined as the fraction of uncertainty in calculated growth rate over the growth rate itself (σ_µ_ / µ) where quantifiable growth rate is where Relative Error = (σ_µ_ / µ) *×* 100 % < 50% (**Fig. 3**).

The value of the water hydrogen assimilation constant (*a_w_*) was determined for both organisms from the slopes of the linear regression relating *^2^F_biomass_* vs. *^2^F_water_*. This value, *a_w_ = X_w_ ×* α*_biomass/w_*, corresponds to the isotopic offset between microbial biomass relative to its growth water. This term includes both *X_w_*, the mole fraction of lipid H sourced from growth water, and α*_biomass/w_*, the isotopic fractionation between whole-cell biomass and growth water (21).

### Nanoscale Secondary Ion Mass Spectrometry

Samples were analyzed with a CAMECA NanoSIMS 50L (CAMECA, Gennevilliers, France) housed in the Caltech Microanalysis Center at the California Institute of Technology. Before, analysis, cells were sputter-coated with 25nm of Au. Cells were analyzed using a 2.5 pA primary Cs+ beam current, and a presputter time of 6 – 20 minutes depending on the size of the raster area. Two masses were collected in parallel (^1^H^-^, ^2^H^-^) using electron multipliers. Individual samples were identified using the NanoSIMS CCD camera and correlated with Raman-measurements using previously generated image maps. For all analyses, at least two frames were collected. All ion images were recorded at 512 x 512-pixel area. Single-cell isotope values were quantified using a custom script (see *Data Availability*) relying on the *sims* Python library (https://github.com/zanpeeters/sims). Cell areas were manually defined using the GNU Image Manipulation Program (GIMP). Isotope ratios (*^2^R*) were converted to fractional abundances (*^2^F*) with the relations: *^2^F = ^2^R / (1 + ^2^R)* and *^2^R = ^2^H / ^1^H,* where ^1^H and ^2^H represent total ion counts in detectors EM1 and EM2, respectively, averaged across a cell area. Additionally, a deadtime correction (70) was applied:

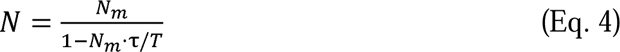

where *N* is the real number of secondary ions, *N_m_* is the number of secondary ions measured, τ is deadtime (in seconds) and *T* is dwell time (seconds per pixel). This instrument’s detector deadtime was 44 ns. Deadtime correction had a negligible effect on isotopic values (**Supplementary Text, Supplementary Fig. S7**).

## Author Contributions

Tristan Caro and Srishti Kashyap designed the study and performed the experiments. Tristan Caro, Srishti Kashyap, Claudia Chen, and George Brown analyzed the data. Alexis Templeton and Sebastian Kopf provided financial support for the study. Tristan Caro and Sebastian Kopf designed the GUI. Alexis Templeton and Sebastian Kopf contributed to the design of the study and interpretation of its findings. Tristan Caro wrote the manuscript with input from all authors.

## Data Availability

Raw data and analysis scripts used to process data and generate visualizations are available at https://github.com/tacaro/Caro-et-al-Raman-SIP. The interactive GUI referenced in this manuscript is available at https://apps.kopflab.org/login with login username *shiny-rSIP* and password *public-access.* In addition, a copy can be downloaded for offline use from the manuscript GitHub repository.

## Acknowledgements

The authors would like to thank the University of Colorado Boulder, Department of Geological Sciences for support of this study. This work was also supported by a NASA Exobiology Program grant to Alexis Templeton (80NSSC21K0489) and an Army Research Office grant to Sebastian Kopf (W911NF2120119 / 78484-LS). The authors sincerely thank Yunbin Guan of the Caltech Microanalysis Center for expertise with nanoSIMS measurements. We thank Eric Ellison of the CU Boulder Raman Microspectroscopy Laboratory (RRID:SCR_019305) for assistance with Raman measurements and data analysis. We extend sincere thanks to John Magyar for helpful discussions during the design of this study. We credit Dan Utter for assistance with nanoSIMS data processing and initial drafting of the nanoSIMS data extraction script. We thank Rebecca Wipfler for assistance with laser microdissection microscopy and coupon etching. Tristan Caro was supported by a National Science Foundation Graduate Research Fellowship and through the BioFrontiers Institute of the University of Colorado Boulder. Claudia Chen and George Brown were supported by the Laboratory for Interdisciplinary Statistical Analysis at the University of Colorado Boulder. The Laboratory for Interdisciplinary Statistical Analysis at the University of Colorado Boulder is supported by National Science Foundation Grant No. 1955109.

