## Supplemental Information for "Single-cell measurement of microbial growth rate with Raman microspectroscopy"

**Running title:** Microbial growth rate measured with Raman spectroscopy.

Tristan A. Caro [a#], Srishti Kashyap [a], George Brown [b], Claudia Chen [b], Sebastian H. Kopf [a], Alexis S. Templeton [a]

1. Department of Geological Sciences, University of Colorado Boulder, Colorado, United States
2. Department of Applied Mathematics, University of Colorado Boulder, Boulder, Colorado, United States

**# Corresponding Author**
Tristan Caro

2200 Colorado Ave
Benson Earth Sciences Building
Rm. 285. UCB 399
Boulder, CO 80309

**This file includes:**

- Supplementary Text
- Supplementary Figures S1 to S8
- Supplementary References

**Other supplementary materials for this manuscript include the following:**

Supplementary File S1: Correlative Dataset of Raman- and nanoSIMS-derived $^2F_{biomass}$ values
Supplementary File S2: R code for running *Shiny rSIP* GUI

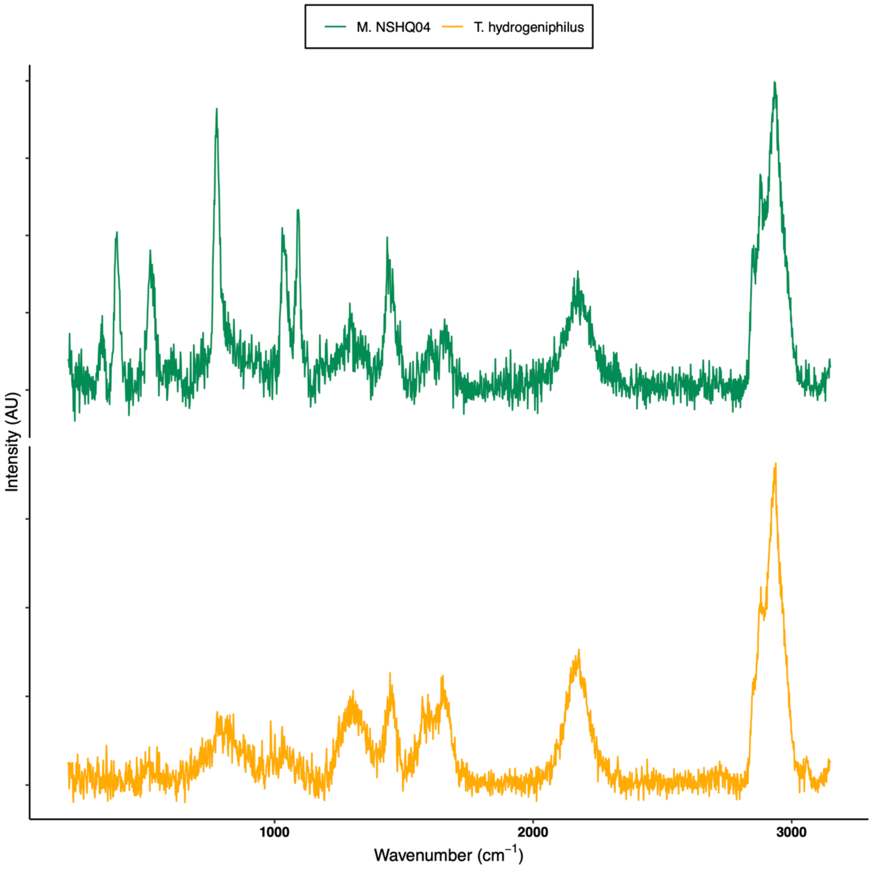


**Supplementary Figure S1.** Representative raw (non-fitted) Raman spectra for organisms *Methanobacterium NSHQ04* (top, green) and *Thermodesulfovibrio hydrogeniphilus* (bottom, yellow) grown at $^{2}F_{L}$ = 50 at. %. Note the unique bands each organism exhibits in the fingerprint region (200 – 1800 cm$^{-1}$), owing to their unique biomolecular makeup.

***Correlation between Raman and nanoSIMS measurements***

Given the observed depression in deuterium abundance measured by nanoSIMS, we sought to understand whether we could correct nanoSIMS D/H values by correlating them with Raman-derived D/H values. With our correlative single-cell dataset, we investigated the extent to which single-cell ${}^{2}F$ measured with both methods agreed (**Supplementary Figure S2)**.

The data derived from Raman spectroscopy was compared with the data from nanoSIMS via simple linear regression in R (1). Simple linear regression was performed across all cell types combined, then with only *T. hydrogeniphilus,* and only *M. NSHQ04*. Cross validation was used to test the predictive power of the linear regression model on data the model was not trained on. The dataset of 351 measurements across both cell types was randomly split into a training set of 189 measurements and a test dataset of 162 measurements using the built-in sample function in R (**Supplementary Fig. S2**). Both the training and test datasets included both cell types. The model formed on the training data was used to predict on the test data and determine the test error. Bootstrap sampling with 1000 replications was performed to generate 95% confidence intervals for the intercept and slope parameters, using all of the available data. Regression diagnostics plots including residuals vs. fitted, QQ-plot, and a plot of Cook's distance are located in the Supplementary Text (**Supplementary Fig. S3**).

We found that strong correlation was demonstrated between Raman and nanoSIMS ${}^{2}F$ (adjusted $R^2$ = 0.8239). Our model tested across both cell types and yielded a residual standard error of 2.49 % on 349 degrees of freedom (F = 1639 on 1 and 349 DF, p < 2.2E-16). Using the model fit on the training set to predict the testing set yielded a root mean squared error (RMSE) of 2.54 %, which represents the difference, on average, between actual and predicted values (Main Text: **Table 1D**). We therefore conclude that correcting nanoSIMS values by conversion to undiluted equivalents is tractable and quantitative. However, this conversion introduces an additional source of uncertainty that, as with applying a single dilution factor, should be accounted for in downstream calculations. As discussed in the Main Text, the lack of parity between nanoSIMS and Raman-derived values reflects dilution of deuterium due to sample washing, and that different treatments during preparation could amount to different degrees of dilution not explained or explored here.

**Commented [SK1]:** Might be helpful to add an additional statement here to clarify that this relationship reflects dilution due to washing, and that different treatments during preparation could amount to different degrees of dilution not explained here.

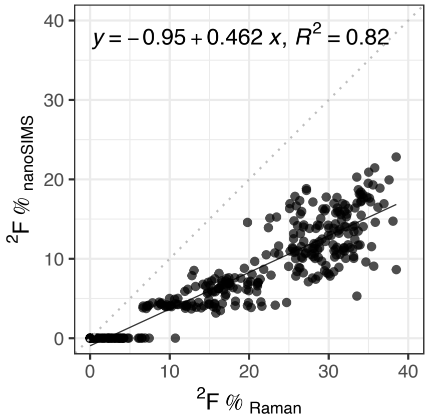


**Supplementary Figure S2.** Correlation between Raman- and nanoSIMS-derived estimates of biomass deuterium enrichment ($^2F$). Each point represents a single cell (n = 351). The deviation from the 1:1 line (dotted line) and scatter around the line of best fit (solid line) are likely due to substantial and variable dilution effects, as discussed in the main text.

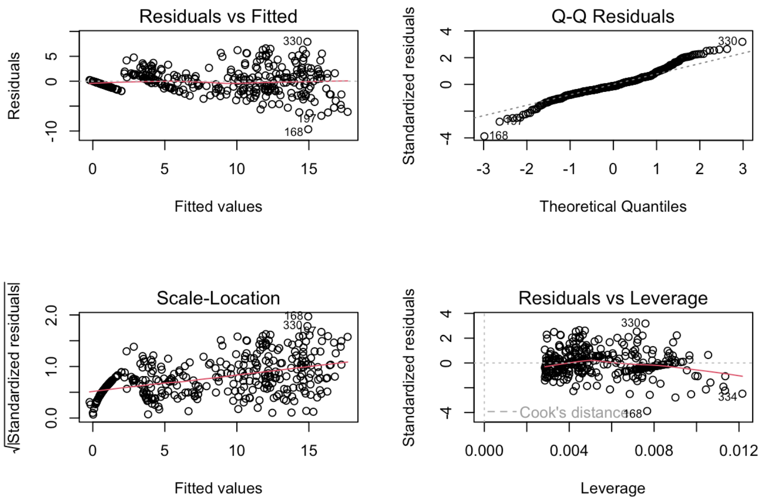


**Supplementary Figure S3.** Model diagnostics plots for simple linear regression of all $^2F_{biomass}$ (*T. hydrogeniphilus* and *M. NSHQ04*) data.

***Surface-Enhanced Raman Spectroscopy (SERS) attempts***

In an effort to achieve higher hydrogen isotopic measurement sensitivity, we attempted to implement surface-enhanced Raman spectroscopy (SERS). For the benefit of future researchers, we report negative results from these efforts.

SERS is a technique that relies on the amplification of Raman scattering of compounds via the adsorption of nanostructured materials, often metal nanoparticles (e.g., Ag, Au, Cu, Zn, etc.), to a target substrate (2–4). Surface enhancement with modern SERS methodologies have resulted in signal enhancements up to $10^5$ - $10^6$ (2). Therefore, we hypothesized that SERS could be a useful technique for increasing the sensitivity of Raman spectroscopy to detect deuterium in microbial biomass.

To this end, we synthesized silver nanoparticles (AgNPs) using the trisodium citrate reduction method adapted from (5–7). We chose AgNPs because this specific SERS substrate has been previously applied for SERS analyses of microbial samples (3,8). AgNPs were synthesized by dissolving 72 mg of AgNO3 in 400mL ultrapure water (18.2 MΩ cm) in an Erlenmeyer flask. The solution was heated to initial boiling (~90˚C) under vigorous stirring. 8 mL of 1% wt/vol trisodium citrate solution was slowly added into the reaction flask and the combined solution was allowed to boil for approximately one hour. Once the solution had turned an apparent luminescent yellow, the solution was allowed to cool to room temperature. The average diameter of AgNPs in suspension was colorimetrically determined to be 50 nm, due to an absorption max was 420nm (9). AgNPs were mixed with sulfate reducing bacteria (SRB) *T. hydrogeniphilus* (See Main Text: Materials and Methods: Biological Sample Preparation). A cell pellet from the SRB culture was mixed directly with 10 µL of concentrated AgNPs and mixed by gentle pipetting, and spotted onto an aluminum-coated glass slide, and allowed to air-dry prior to Raman spectroscopy. Cells were analyzed identically to as described in *Main Text: Materials and Methods* with the following modifications: laser power was held at 10% (0.255 mW) and acquisition time varied between 0.25 and 1 second. Only one acquisition was acquired per cell. For comparison, our non-surface enhanced Raman acquisitions used 10X as much laser power and acquired spectra for 45 seconds, over 2 acquisitions.

**Commented [SK2]:** stimulated is different from SERS

We observed substantial signal enhancement in the fingerprint region on the order of $10^2$ – $10^3$ with the addition of AgNPs. However, we note two key observations that we believe warrant caution when applying SERS methodology to microbial samples. The first, most crucial observation for our study, is the lack of signal enhancement in the high-wavenumber regions corresponding to the organic CH and CD bands (**Supplementary Figure S4**). Amplification of spectral components is the result of the proximity of the adsorbed nanoparticles to specific molecular components (4). We suspect that, because metal NPs preferentially nucleate around highly-functionalized molecules such as fused-ring systems, flavins, nucleic acids, etc. (8), signal enhancement of C-H bonds could be limited due to the lack of interaction between polar nanoparticles and the non-polar regions where many C-H/C-D bonding environments are situated (alkyl chains, fatty acids chains, etc.). The second problem we observed was that signal enhancement of organic features in the fingerprint region (200 – 1800 $cm^{-1}$) was uneven and changed over time with multiple acquisitions, a finding that has been previously reported (8) (**Supplementary Figure S5**). While SERS allows the acquisition of cell spectra in mere seconds, it also produces spurious, ephemeral spectral artifacts, including "ghost peaks" that appeared and disappeared, shifts in peak position, and eventual graphitization ("burning") of the cell. Increased acquisition time led to more rapid graphitization of the cell. Based on our results, we concluded that this SERS methodology, in its current form, is not suitable for measuring cellular deuterium incorporation. More fundamental, mechanistic work is required before it can be robustly applied to environmental systems or to accurately detect and quantify hydrogen isotopic changes.

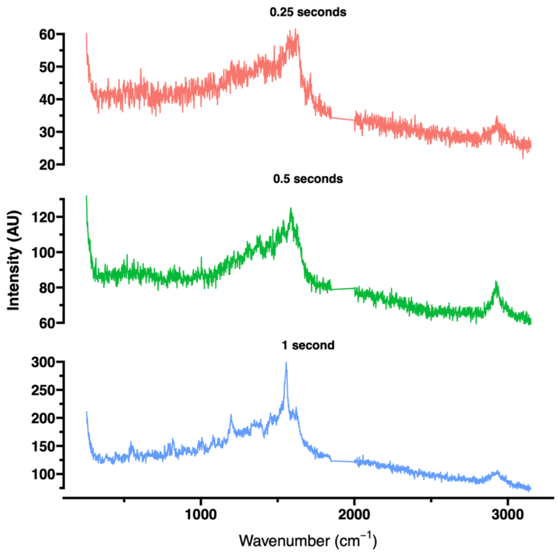


**Supplementary Figure S4.** Representative SERS spectra of *T. hydrogeniphilus* grown in 30% $^{2}H_2O$ across three different acquisition times. Note the highly amplified organic region between 1000 and 1800 $cm^{-1}$ and the poorly-amplified CH region (2800 $cm^{-1}$). The CD band is not observed. The gaps between 1800 and 2000 $cm^{-1}$ represent a spectral region that was not measured for these experiments.

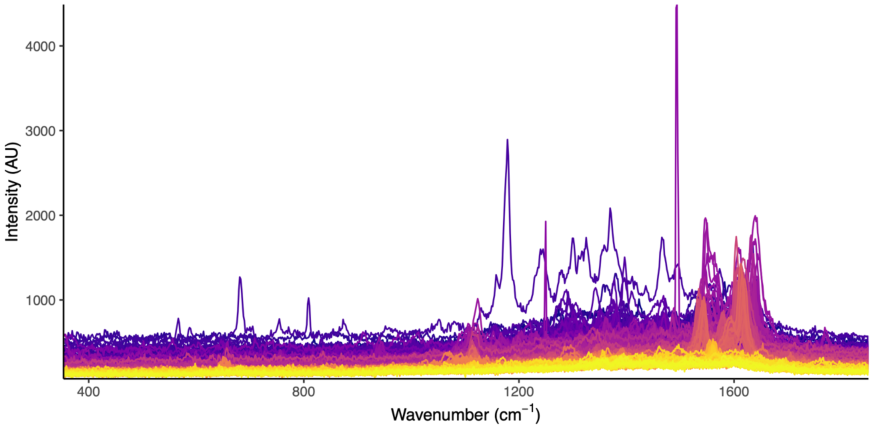


**Supplementary Figure S5.** SERS fingerprint region spectra taken of a single cell across n = 251 independent 0.5 second acquisitions. Line color indicates the timepoint where dark purple is n = 1 and bright yellow is n = 251. Note the rapid shifting of peaks in initial acquisitions and “ghost peaks” appearing and disappearing throughout.

***Quantifying the fraction of exchangeable hydrogen in cell biomass***

After observing the dilution of nanoSIMS-derived ${}^{2}F$, we sought to question the specific mechanism by which nanoSIMS-derived ${}^{2}F$ values became diluted. Specifically, was the observed depression in ${}^{2}F$ primarily the result of exchangeable hydrogen sites that had equilibrated with wash buffers, or were they the result of relict water that remained adsorbed to and within the cell? Using a previously determined molecular inventory of bacterial cells (10) and the molecular structures of dominant cellular biomolecules, we calculated the fraction of hydrogen that is exchangeable in each biomolecule.

**Commented [SK3]:** Do we need to somehow caveat that we're applying this to both bacteria and archaea?

**Commented [TC4R3]:** Added such a caveat to the end of this paragraph.

To estimate the relative contributions of (i) intra/extracellular water and (ii) exchanged biomass hydrogen on nanoSIMS hydrogen isotope measurements, we generated a rough estimate of the cellular hydrogen fraction that is exchangeable with the aqueous environment. We used the molecular inventory published by Neidhardt (10) to estimate the relative mole fractions of different biomolecules. In our calculations, we assumed that hydrogens existing in exchangeable covalently bonded sites (O-H, N-$H_x$, S-H) would rapidly exchange with natural abundance isotopic solutions (11,12). To calculate the mole fraction of hydrogen in a cell that is exchangeable, we need to estimate (i) the total moles of H in a cell and (ii) the fraction of these that are exchangeable. We postulate that this follows a mass-balance relationship such as:

$$F_{H_{ex}} = \frac{H_{ex}}{H_{ex}+H_{non}} \qquad \text{(Eq. S1)}$$

$$H_{total} = H_{ex} + H_{non} \qquad \text{(Eq. S2)}$$

Where $H_{total}$ is the total moles of H in a cell, $H_{ex}$ is the moles of hydrogen in a cell that are bonded at an exchangeable site, $H_{non}$ is the moles of hydrogen in a cell at non-exchangeable sites, and $F_{Hex}$ is the fraction of cellular hydrogen that is exchangeable. From the Neidhardt dataset, we have constraints on the mole fractions of major biomolecules (**Supplementary Figure S6A**) as well as their proportions relative to the dry weight of the cell. We examined the molecular structures of the biomolecules within this inventory and manually tabulated whether hydrogens in these structures existed at exchangeable sites (O-H, N-$H_x$, S-H) (**Supplementary Figures S6B**). We calculated that, per g of cell dry weight, there exists 62944.4 µmol of hydrogen, out of which 13483.6 µmol exists at exchangeable sites. This corresponds to a 21.4 % fraction of exchangeable H. Proteins contributed the highest quantities of exchangeable H due to the high functionalization of peptide residues, whereas lipids contributed very little (**Supplementary Figure S6B**).

Given our observed dilution factor of 58.1% and our estimation that roughly one-fifth of bacterial H is readily exchangeable, our calculations suggest that there exists a contribution from relict intracellular and/or extracellular natural-abundance water to total measured hydrogen isotopic composition. Drying of cells on a slide and placing samples under vacuum in the nanoSIMS instrument may not sufficiently desiccate cells for un-biased hydrogen isotopic analysis. We emphasize that this is a rough estimation based on a single cultured representative and that the cellular contents of microorganisms can substantially differ. More work is needed to generate molecular inventories of cells outside of common cultured representatives and across a broader range of phylogeny.

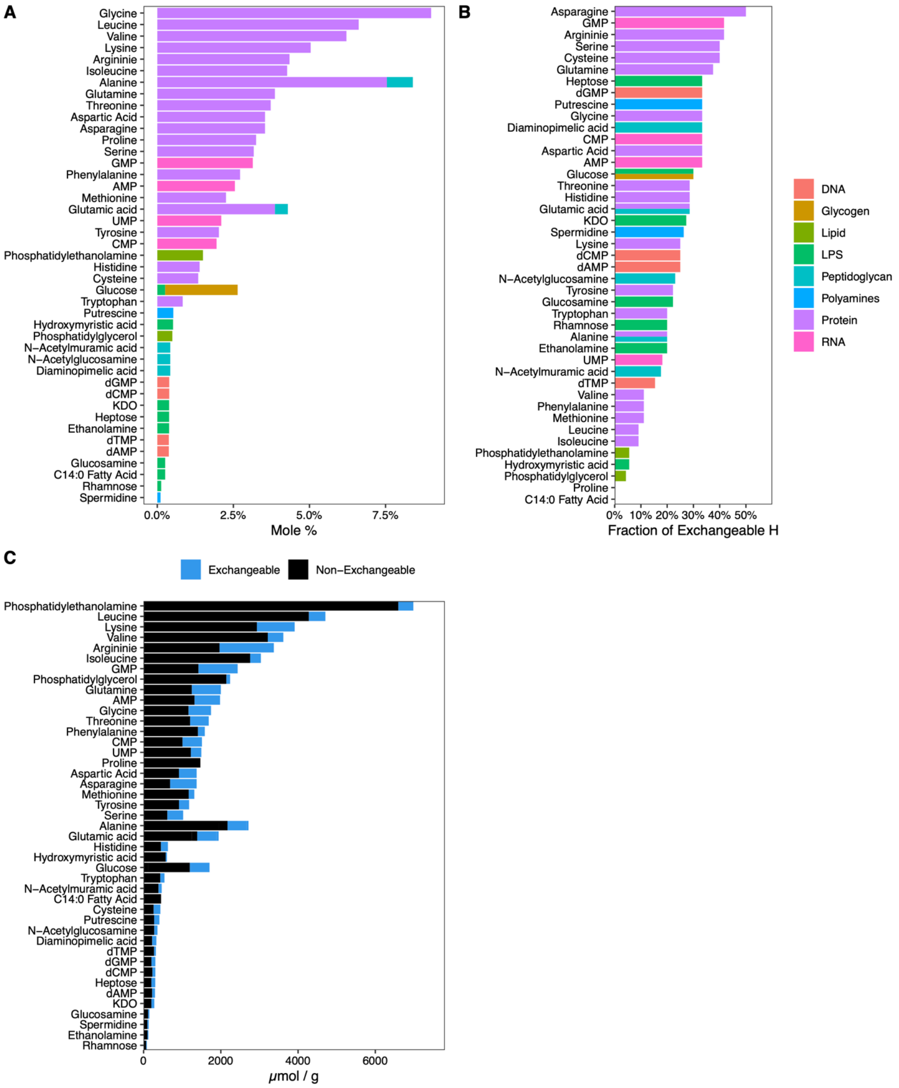


**Supplementary Figure S6: (Panel A)** We calculated the relative mole percent of each major biomolecule in the molecular inventory (10). **(Panel B)** Using the molecular structures of each biomolecule, we determined the fraction of component hydrogen that exists at exchangeable bonding sites. **(Panel C)** The total μmol per gram (dry weight) of a generic bacterial cell that consists of exchangeable (blue) and non-exchangeable (black) hydrogen atoms.

***NanoSIMS Dilution Factor Calculations***

Dilution of isotopic signal by sample preparation for nanoSIMS has been widely observed (13–17). Here we define the dilution factor of sample preparation (D) similar to (18):

$$D = \frac{F_{after} - F_{before}}{F_{added} - F_{before}} \quad \text{(Eq. S3)}$$

where *F* denotes isotope fraction (at. %) of biomass before ($F_{before}$) and after ($F_{after}$) sample preparation, as well as that of the diluent material ($F_{added}$). Here, we take the Raman-derived $^{2}F$ as the *before* value, the nanoSIMS as the *after* value, and natural abundance hydrogen as the *added* value. Because the natural abundance of deuterium is negligible in comparison to the label strengths used for this study, we set $F_{added}$ = 0.00015 % (VSMOW). Here we make explicit the assumption that the Raman-derived value, calculated from CD%, is approximate to whole-cell isotopic composition *before* cell washing. In other words, we assume that molecules incorporate deuterium equally towards non-exchangeable C-H bonds (measurable by Raman) as protic sites in cellular biomass (measurable by nanoSIMS).

We calculated uncertainty in dilution factor using standard propagation of uncertainty via partial derivatives, assuming all measurements to be uncorrelated. This uncertainty term is calculated to be:

$$\sigma_D = \sqrt{\left(\frac{1}{F_{added} - F_{before}} \cdot \sigma_{F_{after}}\right)^2 + \left(\frac{F_{after} - F_{added}}{(F_{added} - B)^2} \cdot \sigma_{F_{before}}\right)^2 + \left(\frac{F_{before} - F_{after}}{(F_{added} - F_{before})^2} \cdot \sigma_{F_{added}}\right)^2} \quad \text{(Eq. S4)}$$

For future researchers: back-calculation of the isotopic composition of biomass *before* sample preparation can be done by rearranging Eq. S3, solving for $F_{before}$:

$$F_{before} = \frac{F_{after} - (F_{added} \cdot DF)}{1 - DF} \quad \text{(Eq. S5)}$$

The propagated uncertainty in this equation is therefore:

$$\sigma_{F_{before}} = \sqrt{\left(\frac{1}{1 - D} \cdot \sigma_{F_{after}}\right)^2 + \left(-\frac{D}{1 - D} \cdot \sigma_{F_{added}}\right)^2 + \left(\frac{F_{after} - F_{added}}{(1 - D)^2} \cdot \sigma_D\right)^2} \quad \text{(Eq. S6)}$$

***Effect of dead time correction on nanoSIMS-derived data***

For our nanoSIMS dataset, we applied a detector dead time correction (DTC). Deadtime is a time interval during which an electron multiplier (EM) detector is unable to capture additional signal while the previous signal is being processed. When a secondary ion ($1H^-$, $2H^-$) hits the conversion dynode, it ejects an electron that ejects further electrons across subsequent dynodes in the EM. This results in an electron pulse at the pre-amplifier electronics. During the time it takes to record this signal, the EM cannot measure additional incoming ions arriving at the conversion dynode. This is problematic when the secondary ion count rate is high as it would effectively depress the recorded signal (19).

We observed a negligible effect of dead time correction (Main Text: *Materials and Methods*) on deuterium isotopic abundances. Our nanoSIMS instrument used the default value of 44 ns deadtime. The maximum deadtime effect was observed at higher deuterium abundance (**Supplementary Figure S7A, B**). The maximum residual difference between raw and dead-time corrected (DTC) nanoSIMS data was just under 0.05 at. %, with most residuals below 0.03 at. %. Given that our labeling enrichment varied between 0 - 50 at. %, we consider this a negligible effect for the purposes of our study **(Supplementary Figure S7C)**.

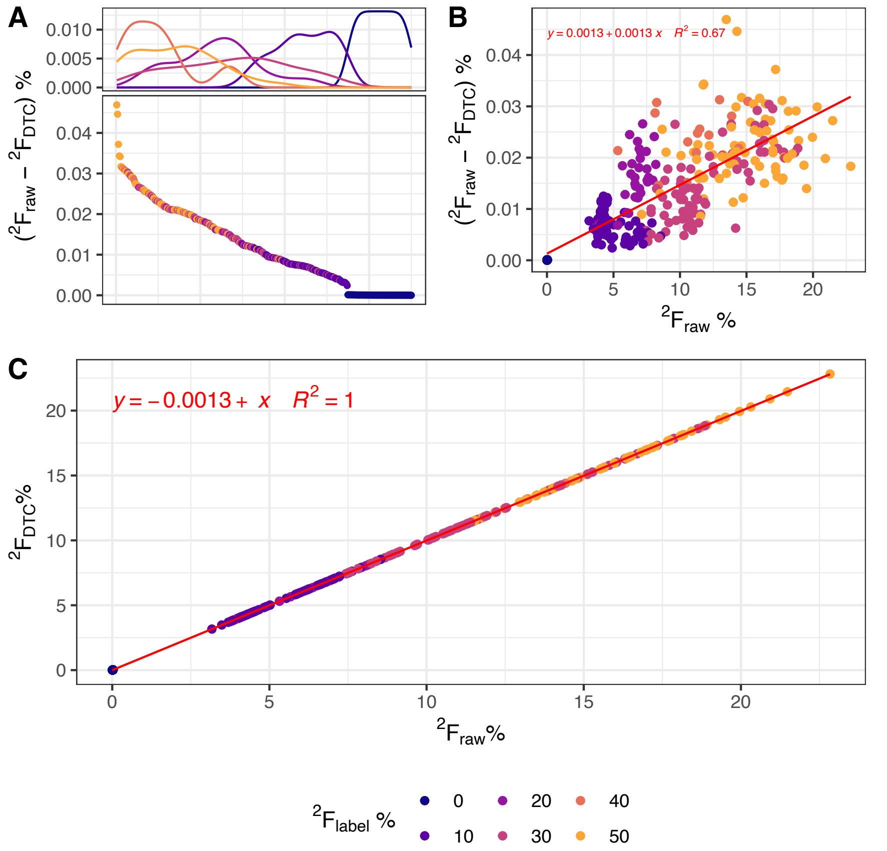


**Supplementary Figure S7.** Effects of dead-time correction on reported nanoSIMS $^2$F values. (**Panel A**) The difference in $^2$F between raw and dead time-corrected (DTC) values. Points are ordered on the x-axis in descending order from greatest residual difference between raw and DTC value. The top panel indicates relative frequency of observations (y-axis), color differentiated by $^2F_L$. (**Panel B**) The difference in $^2F$ between raw and DTC values increases with increasing $^2F$. Residual differences in raw and DTC values are negligible compared to the magnitude of the $^2F$ values reported here. $^2F$ values near natural-abundance exhibited miniscule effects related to dead time correction. (**Panel C**) Raw and DTC $^2F$ values plotted against each other, with a linear fit (red).

**Commented [SK5]:** Can you label the x on this panel as relative frequency?

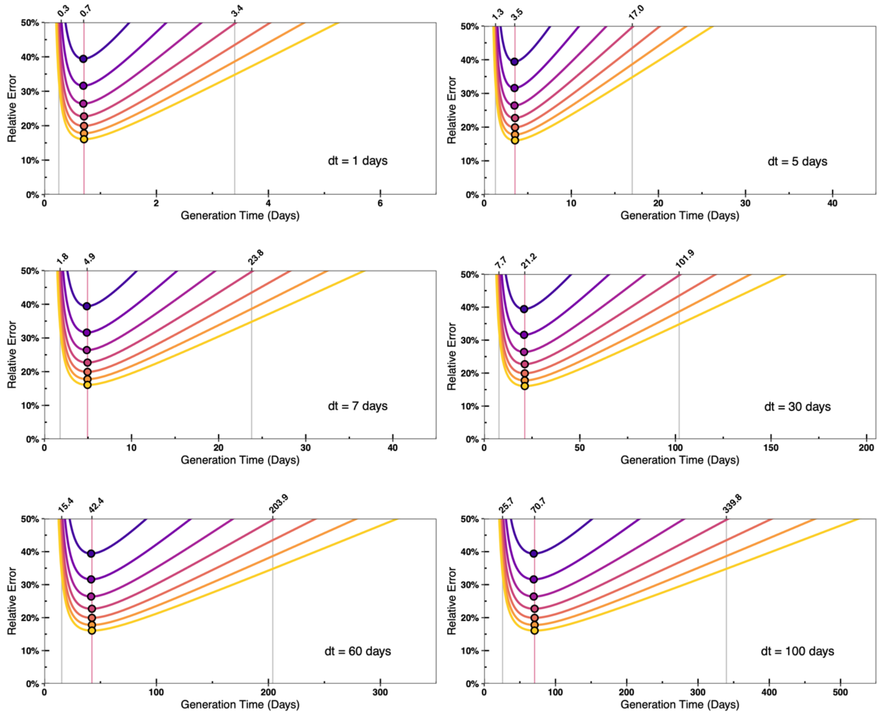


**Supplementary Figure S8.** Ranges of generation time quantification across multiple incubation times (dt). As incubation time increases, longer the generation times can be captured. Similarly, increasing incubation time also increases the minimum generation time that can be captured. Note that the x-axes are scaled differently in each plot. Additional SIP conditions (incubation times, label strengths, etc.) can be examined with our microbial growth model and with the Shiny-rSIP GUI (Supplementary Data).

**Commented [SK6]:** You can state here again that such variable incubation time plots can be determined using the shiny app.
